# Nutrition or nature: disentangling the complex forces shaping prokaryote pan-genomes

**DOI:** 10.1101/2020.12.14.422685

**Authors:** Daniel R. Garza, F. A. Bastiaan von Meijenfeldt, Bram van Dijk, Annemarie Boleij, Martijn A. Huynen, Bas E. Dutilh

## Abstract

Microbial pan-genomes are shaped by a complex combination of stochastic and deterministic forces. Even closely related genomes often exhibit extensive variation in their gene content. Understanding what drives this variation requires exploring the interactions of genes with each other and with their external environments. However, to date, conceptual models of pan-genome dynamics often represent genes as independent units and provide limited information about their mechanistic interactions. Here, we use pan-reactomes as proxies for pan-genomes since they can explicitly represent the interactions between the genes that code for metabolic reactions and simulate complex phenotypes that interact with the metabolic environment. We interpreted pan-reactomes as dynamic pools of metabolic reactions that are potentially gained or lost and simulated the routes along which different lineages lose reactions in alternative environments. We performed these simulations on the pan-reactomes of 46 bacterial and archaeal families covering a broad taxonomic range. These simulations allowed us to disentangle metabolic reactions whose presence does, and does not depend on the metabolite composition of the external environment, allowing us to identify reactions constrained “by nutrition” and “by nature”, respectively. By comparing the frequency of reactions from the first group with their observed frequencies in bacterial and archaeal families, we predicted the metabolic niches that shaped the genomic composition of these lineages in their evolutionary past. Moreover, we found that the lineages that were shaped by a more diverse metabolic niche also occur in more diverse biomes as assessed by global environmental sequencing datasets. Together, we introduce a computational framework for analyzing and interpreting pan-reactomes that provides new insights into the ecological and evolutionary drivers of pan-genome composition.

## Introduction

In the evolution of microbial genomes, genes are gained and lost by means of mutations, insertions, deletions, duplications, and horizontal gene transfers (HGTs) (Lorenz and Wackernagel, 1994; Paget and Simonet, 1994; Puigbò et al., 2014; Thomas and Nielsen, 2005). As a result of these processes, gene content varies significantly even between closely related genomes (Hao and Golding, 2006; Iranzo et al., 2019; Puigbò et al., 2014). The long-term outcome of genome evolution events—measured by the frequency of genes in the pan-genomes of microbial lineages—is determined by a combination of stochastic and deterministic factors. For instance, if two genes code for similar functions and there is no effective fitness difference, the loss of one over the other would likely be a stochastic event. Conversely, if there is a difference in fitness effect between the encoded functions due to the microbe’s physicochemical environment, the loss of one would likely be a deterministic event. Because of the complex structure of the molecular network in a cell, it is challenging to disentangle the forces that shape the gene content of genomes, a necessary step for the development of theoretical frameworks to interpret the gene frequency of pan-genomes.

Recent studies have applied the results from population genetics theory—based on using the frequency of gene variants (genotypes) to study the evolutionary phenomena that shape populations—to explain patterns in the gene frequency of microbial pan-genomes (McInerney et al., 2017; Rocha, 2018; Sela et al., 2016; Shapiro, 2017). This has been justified since pan-genomes, instead of populations, can be viewed as the key units of prokaryote evolution (Puigbò et al., 2014) and the frequency of genes in a pan-genome can be translated to the frequency of genotypes. There are caveats in the immediate translation of traditional population genetics theory to pan-genomes since the frequency of genotypes and the frequency of genes reflect different phenomena. While the former reflects selection coefficients, the mutation rates, and effective population sizes, the latter reflects HGT rates and genomic adaptation to the environment (Aminov, 2011; Bonham et al., 2017; Hao and Golding, 2006; Maistrenko et al., 2020; Sitaraman, 2018; Wiedenbeck and Cohan, 2011). These caveats can be addressed by developing more realistic frameworks to model and explain gene frequency in pan-genomes.

The genes in the pan-genome of a microbial clade are more accessible to the strains of this clade than of other clades. In general, the probability that a recipient microbial genome will integrate foreign DNA increases exponentially with an increase in the similarity between the donor DNA and the recipient chromosome (Dixit et al., 2017). As a result, closely related genomes share more genes than distantly related ones. This asymmetry in the distribution of genes is an important factor to consider in order to develop realistic explanatory models for the gene frequency of pan-genomes.

Another important factor to consider is that the acquisition of new genes by genomes is counterbalanced by the frequent loss of genes (Bolotin and Hershberg, 2016; Puigbò et al., 2014; Snel et al., 2002; Wolf and Koonin, 2013). Gene loss is majorly a clock-like process, where genes under weak or no selection tend to be inactivated by random mutation and lost by deletion (Bolotin and Hershberg, 2016; Sheinman et al., 2020; Wolf et al., 2012). This process is widely observed across microbial genomes and virtually all species with genomes smaller than 2 Mb evolved from ancestors with substantially larger genomes (Makarova and Koonin, 2007; Ochman, 2005; Wolf et al., 2012). Gene loss is also the major source of genomic variation of intracellular parasites that do not undergo extensive HGT (Moran and Mira, 2001) and of bacteria that are adapted to stable and nutrient-rich environments, such as host-associated microbiomes (Makarova and Koonin, 2007). Based on the high rates of gene loss, genes that are not under selective pressure may eventually be lost.

The frequency distributions of genes fit mathematical functions with regular and universal shapes (Koonin, 2011; Mazzolini et al., 2018; Pang and Maslov, 2013). One example is the asymmetric U-shape that is observed for a broad range of prokaryote groups (Koonin and Wolf, 2008; Touchon et al., 2009). This distribution exhibits a pronounced number of core genes to the right (frequency approx. 1), a spread of moderately common genes in between, and a peak of rare genes to the left (frequency << 1). Frequency is often confounded with the relative essentiality of a gene and said to reflect its adaptive value in the face of purifying selection. In this view, core genes are essential under any condition, while increasingly rare genes are increasingly dispensable. Such propositions were challenged by simple neutral models—that do not attribute different selective advantages to different genes—that reproduced the characteristic U-shape and exhibited a reasonable fit with the distribution of genes across the genomes of several clades (Baumdicker et al., 2012; Collins and Higgs, 2012; Haegeman and Weitz, 2012). But non-neutral models can also exhibit a good fit to the characteristic U-shape distributions. A model that attributes varying selective advantages for genes was found to be a better fit to several pan-genomes compared to the neutral alternatives (Lobkovsky et al., 2013).

The simplified ‘bag-of-genes’ models described above do not explicitly consider gene functions and their interactions. These models provide important insights into the evolutionary dynamics of pan-genomes, but ignore the functional forces driving gene frequencies and, more importantly, do not provide a mechanistic interpretation for the variation in gene content. In nature, selection acts on the phenotype, and microbes exhibit complex phenotypes that result from the combined action of multiple gene products. In many cases, phenotypes are dependent on the interactions of gene products with the environment.

A promising approach to integrate the functions and interactions of genes into models of genome evolution is to use the genome-encoded metabolic reactions, the reactome, as a proxy for the gene content of genomes. Reactions from the reactome can be integrated into a functional network (also referred to as a genome-scale metabolic model (GSMM)) that represents the genotype-to-phenotype map (Edwards and Palsson, 1999; Gu et al., 2019; Norsigian et al., 2020; O’Brien et al., 2015). The molecular components (protein-encoding genes) of reactomes are readily inferred from microbial genome sequences (Gu et al., 2019; Hosseini et al., 2016; Wagner, 2012). Similar to pan-genomes (see reviews (Lapierre and Gogarten, 2009; Medini et al., 2008, 2005)), pan-reactomes can be subdivided into the core-reactome, consisting of reactions present in all strains, and the flexible reactome, consisting of reactions found in at least one or a few strains. Functional networks derived from pan-reactomes are also capable of simulating complex phenotypes, such as the conversion of energy and matter from diverse environmental metabolites into sugars, nucleic acids, lipids, and proteins (Gu et al., 2019).

Here we used pan-reactomes as models to simulate and understand patterns in pan-genomes. In order to incorporate realistic features of pan-genome evolution, we used pan-reactomes as proxies for pools of genes that are accessible to related strains by HGT. We then modeled alternative routes of gene loss by sampling minimal functional reactions sets in diverse environment compositions. These minimal functional reaction sets have similar properties as the previously defined elementary flux modes (EFMs) used to identify functional pathways in reactomes (Zanghellini et al., 2013), thus, we termed our sets panEFMS (pan-reactome elementary flux modes) and used them to distinguish two important drivers of reaction frequencies in pan-reactomes, which we refer to as ‘nutrition’ and ‘nature’. The frequency of reactions that are driven by nutrition depend on the environment composition, while the frequency of reactions that are driven by nature does not depend on the environment composition. Our framework mechanistically disentangles environment-driven from environment-independent reactions and uses their distribution in panEFMs to build a model that predicts the metabolite preferences of pan-reactomes from their environment-driven reactions. We applied this model to the pan-reactomes of 46 bacterial and archaeal families, allowing us to assess the patterns of microbial genome evolution that result from the function and interaction between metabolic genes.

## Results

### Functional reactomes of individual strains are samples from a pan-reactome

Our model consists of directed bipartite graphs of reactions and metabolites (Figure 1A) derived from reactomes of related organisms (Figure 1B) that together form a pan-reactome (Figure 1C). To illustrate the model in a tractable way, we use a toy model to illustrate how environment-independent and environment-driven processes together shape the frequency distribution of reactions in pan-reactomes (Figure 1, Table S1). We expand the example by applying the model to the pan-reactome of the *Aeromonadaceae* family that was built based on the metabolic reactions encoded in the genomes of 135 strains that belong to different *Aeromonadaceae* species (Table S2). This pan-reactome network contains 1796 reactions and 292 environment compounds (Table S3). These reactomes take up compounds from the external environment (MX_e metabolites in the toy model in Figure 1) and convert them to synthesize metabolites required for biomass production (M10_i, M11_i, and M12_i). As described in the Introduction, evolutionary processes sample subsets of reactions from this pan-reactome to generate reactomes of individual strains. Strain reactomes are considered functional if they can synthesize all the biomass compounds and do not accumulate by-products, like metabolite M2 in the first reactome of Figure 1B that needs to be exported by reaction R10. Three examples of viable reactomes are shown in Figure 1B and others can be formed. For the family *Aeromonadaceae*, significantly more viable reactomes can be formed than those of the 135 sequenced strains (Table S2), which together defined the family-level pan-reactome. Thus, a pan-reactome defines a space of potential functional reactomes, some of which are realized by actual strains in nature.

**Figure 1:**
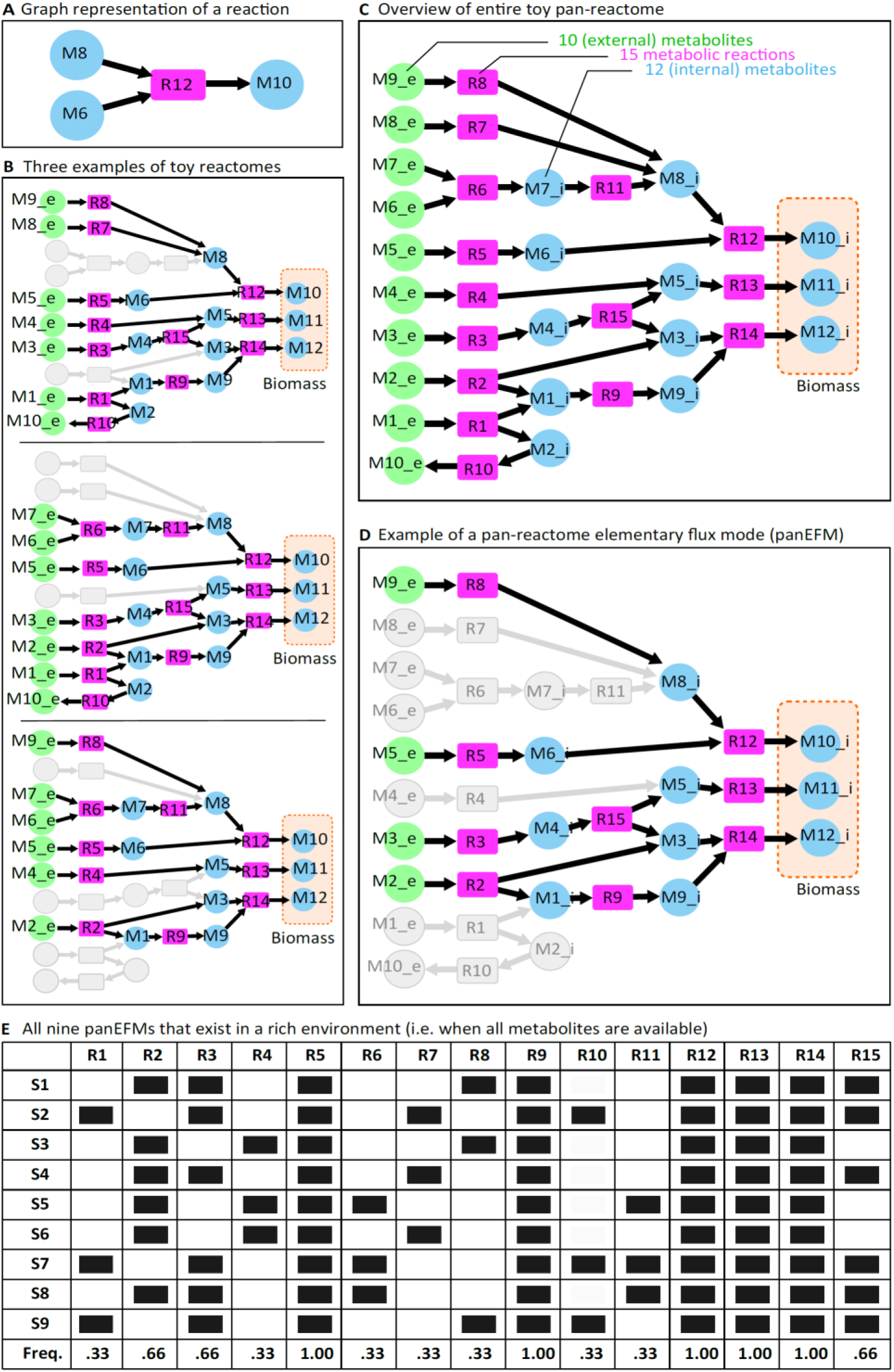
Toy model. (A) Example of a metabolic reaction. Reactants and products are depicted as circles and reactions as rectangles. R eaction directionality is indicated by the arrows. (B) Three functional reactomes derived from the toy model, each capable of synthesizing the biomass compounds M10_i, M11_i, and M12_i from the environmental precursors (‘MX_e’ compounds depicted with green circles). (C) Pan-reactome network aggregating reactions from the different reactomes into a single network. The “_e” and “_i” termination of metabolites denote internal and external metabolites, respectively. (D) An example of a panEFM. Each reaction in this network is essential since its removal would impair the synthesis of the biomass components. (E) Collection of all nine possible panEFMs that can be created from the reactions in this toy pan-reactome in a rich environment. Dark squares denote the presence of reactions. The frequency of reactions across the collection of panEFMs is shown in the last row.

While evolutionary processes constrain the functional reactomes that can be sampled from a pan-reactome pool, the reactions that are selected in practice depend on the environment (environment-driven reactions, ‘nutrition’) and on the network (environment-independent reactions, ‘nature’). Using the toy model as an example, in Figure 1C reactions R5, R9, R12, R13, and R14 are environment-independent reactions that are always required for biomass production, they are irreplaceable in the synthesis of essential biomass precursors and are thus necessarily present in each functional reactome. In complex natural networks, some environment-independent reactions may also depend on the presence of other reactions in the network, as we will see below. In contrast, the network may synthesize the biomass precursor M8_i by either using reactions R8, R7, or a combination of R6 and R11. In principle these three metabolic routes are equivalent, but the presence of external metabolites determines which ones are functional and these reactions are considered environment-driven.

### Pan-reactome elementary flux modes (panEFMs) predict reaction frequencies in the pan-reactome

To explore the space of possible reactomes within a group of organisms that comprise a pan-reactome, we modelled functional reactomes under the hypothesis that evolution tends to lose non-essential reactions, by sampling panEFMs from the pan-reactome. A single panEFM consists of a group of reactions that together are functional in a defined environment, but the removal of any reaction makes the network non-functional. An example of a panEFM is shown in Figure 1D (set S1). The number and composition of panEFMs depends on the specific environment, with rich environments having more panEFMs. For example, our toy model has nine panEFMs (sets S1-9) in a rich environment where all external metabolites are available. The panEFMs from the pan-reactome allow us to generate an expected frequency of each reaction in the pan-reactome in the context of a defined metabolic environment. For example, given a rich environment the bottom row of Figure 1E shows the expected frequency of reactions in the lineage represented by the toy pan-reactome, given that in this model there is no selective advantage of using one pathway over another to synthesize a specific metabolite.

Due to the combinatorial explosion involved in extracting all possible panEFMs from large pan-reactomes, for bacterial and archaeal families we used a random sampling approach to approximate the space of all possible panEFMs. We first determined that fewer than 200 panEFMs sampled on different random environments were sufficient for convergence to an average reaction frequency distribution with 99% reproducibility and a mean-squared error approaching zero (Figure S1). We thus generated one million panEFMs for each family, including 1000 panEFMs sampled in each of 1000 different random environments (see Methods).

### Disentangling the forces driving the frequency of reactions in pan-reactomes

To tease apart the forces that shape pan-reactome composition and infer to what extent the evolution of each reaction is driven by nature versus nutrition, we calculated an environment-driven score (EDS) that ranges from zero (environment-independent, frequency not dependent on the metabolic environment, nature) to one (environment-driven, frequency dependent on the metabolic environment, nutrition). First, we calculated the absolute difference between the predicted reaction frequency in one specific virtual environment and its mean frequency across all 1000 random virtual environments (see Methods, cf. last row in Figure 1E), i.e. the residual. Because the latter value reflects the mean frequency overall, this residual quantifies the extent to which the frequency of a reaction is different in each specific environment. We calculated the EDS for each reaction as the scaled standard deviation of these residuals, as this reflects to what extent the reaction varies across specific environments. We illustrate the EDS score in the toy model (Figure 2A) where reactions R5, R9, R12, R13, and R14 are identified as environment-independent (EDS = 0.0), while the rest are environment-driven (EDS > 0). We used this approach to estimate EDSs for all reactions across all bacterial and archaeal families (Table S4). For the *Aeromonadaceae* family, we found that the top environment-driven reactions are reactions involved in the degradation of valine, leucine, and isoleucine (Table S4).

**Figure 2:**
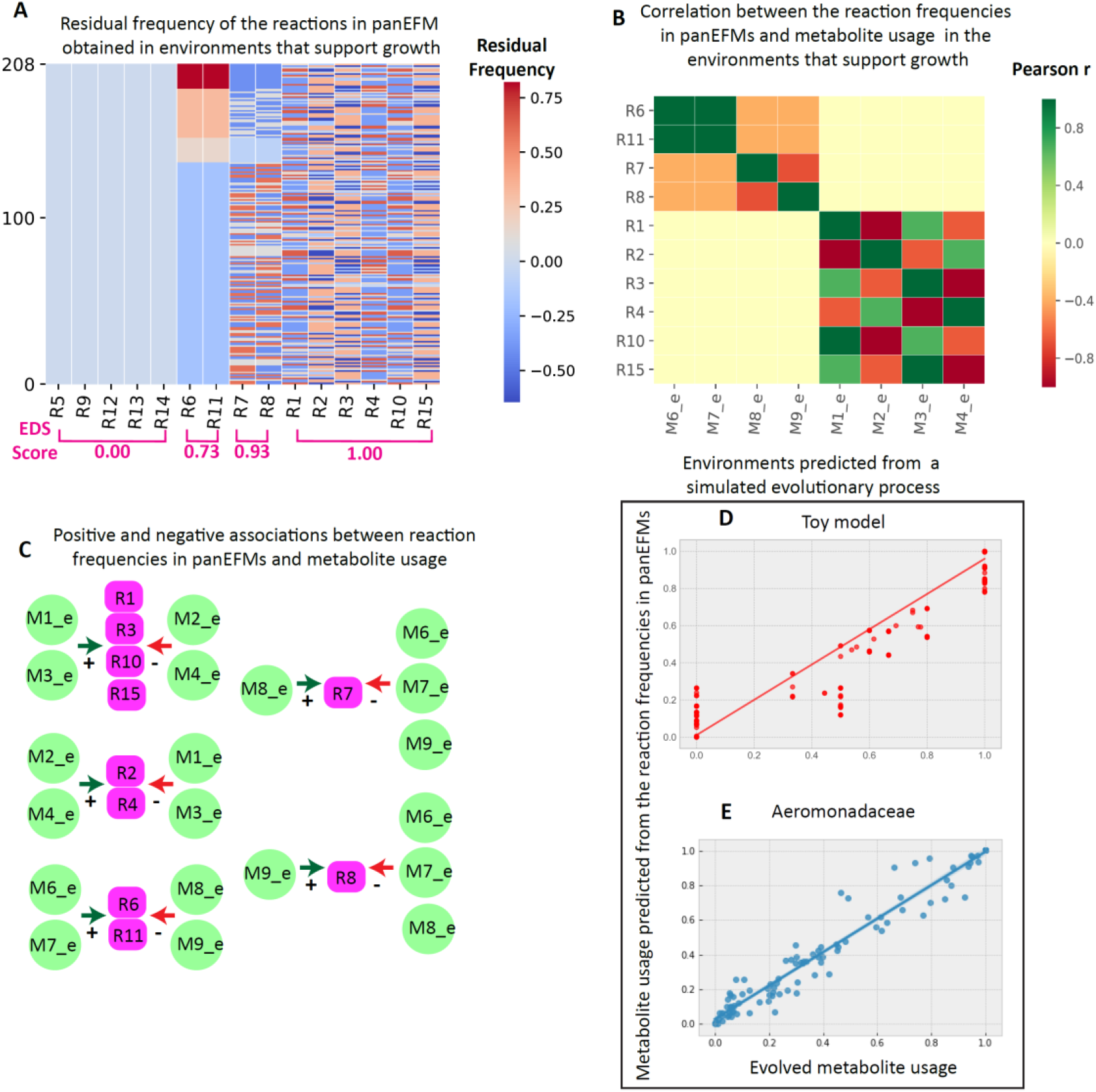
Identifying environment-reaction associations. (A) Residual reaction frequencies predicted from the collection of panEFMs that exist across the 208 environments that support the growth of the toy pan-reactome (Figure 1C). The residuals are the difference between the average frequency over the collection of panEFMs defined within each environment and the average across all environments. Reactions along the x-axis are sorted by the environment-driven score (EDS), see text for details; (B) Metabolite-reaction association matrix defined by the pairwise correlation between the metabolites and reactions with non-zeros residuals; (C) Subnetworks generated from the rows of the metabolite-reaction association matrix (shown in B), showing the positive (+) and negative (−) associations between metabolite usage and reaction frequencies in panEFMs; (D), (E) Elastic net prediction of the metabolite usage that evolved in a simulation of a Moran-like process. The evolved reaction frequencies were used to predict how the resulting strains use metabolites in their environments (y-axis) and compared to their usage in the simulated environment (x-axis).

### Associating environment-driven reactions to environmental metabolites

The EDS score identifies reactions that are environment-driven. Next, we focused on identifying the specific metabolites that drive these frequencies. Because the frequency of reactions in a pan-reactome depends on environmental metabolites in complex ways, we correlated the predicted reaction frequency across 1000 virtual environments to the metabolite usage frequency across those same environments (see Methods). This association, illustrated in Figures 2B and 2C for the toy model, quantifies how external metabolite usage explains the frequency distribution of reactions in pan-reactomes. For instance, the frequency of reaction R8 that produces the biomass precursor M8_i is positively associated with the usage of metabolite M9_e (Figure 2C) and negatively associated with the usage of the metabolites that enable alternative routes to produce M8_i (i.e. M6_e, M7_e, and M8_e; Figure 1C). Note that the frequencies of reactions in the pan-reactome may thus reveal metabolite availability in the environment where the lineage evolved.

### Predicting metabolite usage preferences from reaction frequencies in pan-reactomes

Reaction frequencies may be readily observed in natural pan-reactomes by comparative genomics. To predict the metabolite preferences of prokaryotic families from these reaction frequencies, we trained an elastic net (EN) model on the reaction frequencies in the collection of panEFMs to predict their metabolite usage profiles across the growth-supporting virtual environments (see sections “Reaction frequencies” and “Elastic net” in Methods). We confirmed the accuracy of the EN model for the toy model and the *Aeromonadeceae* pan-reactome by predicting the metabolic niche of reactomes whose evolution was simulated in a defined environment using a Moran-like process of gain and loss of genes (Moran, 1958) (see “Toy model” in the Methods). The EN model accurately predicted the metabolite usage of resulting linages of both the toy model (Figures 2D, r=0.98, p<e-10) and the *Aeromonadaceae* pan-reactome (Figures 2E, r=0.98, p<e-71). Thus, we were confident that we could use the EN to predict the metabolic niche where a pan-reactome evolved based on the extant frequencies of its environment-driven reactions. As described above, these environment-driven reactions are identified by sampling panEFMs across many different environments, so the metabolite usage cannot be directly inferred from the extant reactomes (networks in Figure 1B or the 135 *Aeromonadaceae* GSMMs, Table S2) but requires the intermediate step of sampling panEFMs. We were able to compare evolved metabolite usage with the metabolite usage predicted by the EN because we simulated the evolutionary process in pre-defined environments and could then compute how the evolved reactomes utilize metabolites in these environments. Thus, we proposed an innovative approach to address the elusive question of the preferred metabolic niche of a microbial lineage from the reaction frequencies in its pan-reactome, which in turn can be readily inferred from genome sequences of related strains.

### panEFMs delimit the space of possible pan-reactomes

In the following sections, we will use the framework illustrated above for the toy model and the *Aeromonadaceae* pan-reactome to analyze the pan-reactomes of 46 prokaryote families, each containing more than 24 sequenced genomes (Table S2, see “Pan-reactomes” in Methods).

Reaction frequencies of the collection of panEFMs obtained across random simulated environments reflect an evolutionary landscape of reactomes that could be derived from the family-level pan-reactome pools. Notably, we observed that this landscape exhibited clear family-specific clusters when projected in two dimensions (Figure 3A). The pan-reactomes derived from the sequenced genomes in a family (see “Reaction frequencies” in Methods) were generally found to lie within these clusters (large points in Figure 3A). Thus, our approach of sampling a stochastic distribution of pan-reactomes represented the observed (realized) pan-reactome within this evolutionary landscape.

**Figure 3:**
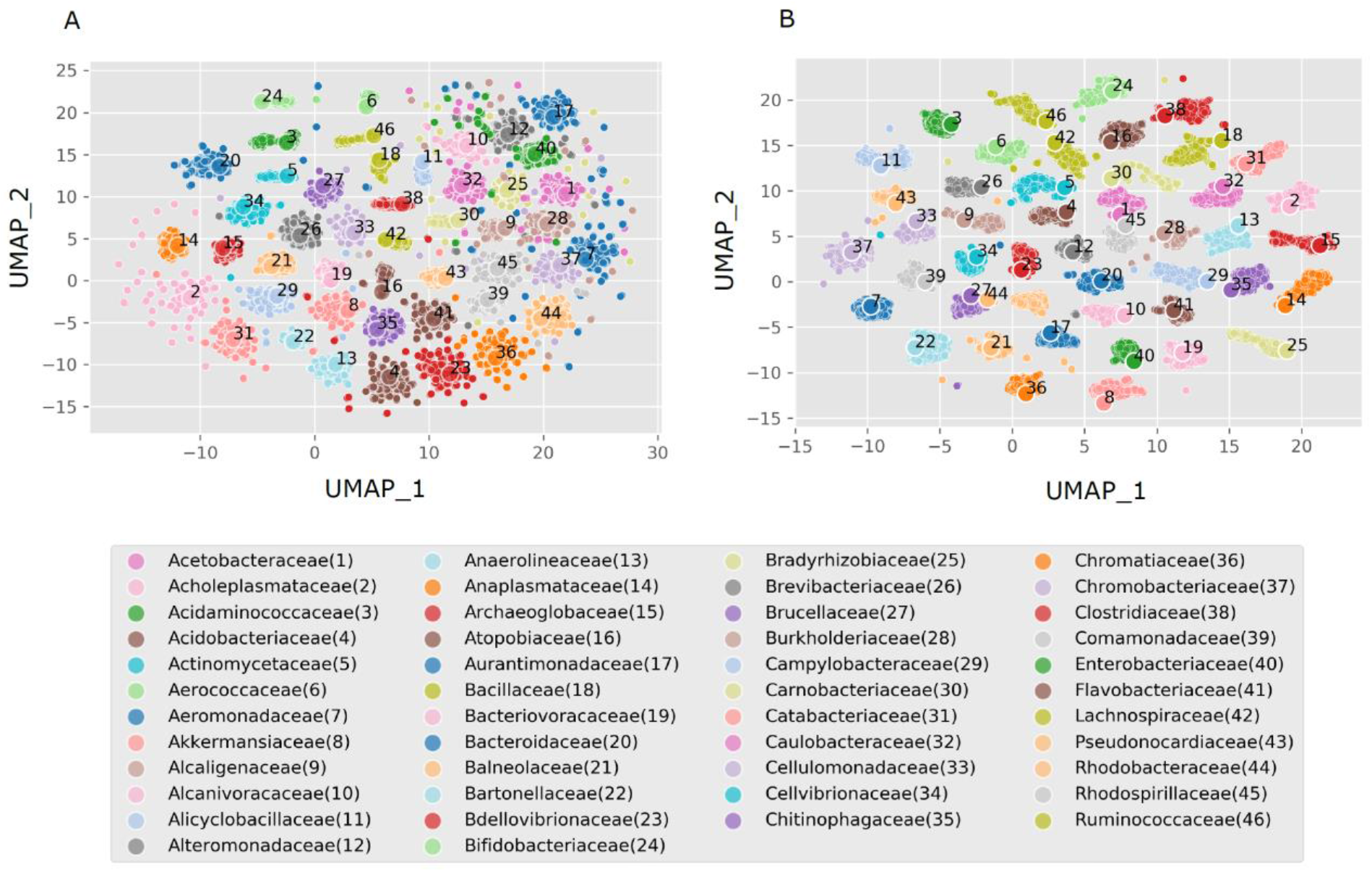
Evolutionary landscape of possible pan-reactome reaction frequencies and metabolite usage profiles based on sampling panEFMs in 1000 random environments. (A) UMAP projection of reaction frequencies in the collection of panEFMs sampled from different prokaryotic families (Table S2). Each smaller point represents the reaction frequency distribution calculated from 1000 panEFMs sampled on one random environment. The large dots are the frequencies observed in the natural pan-reactomes. (B) UMAP projection of the metabolite usage profiles obtained from the same panEFMs projected in A. The large dots are the elastic net (EN) predictions of these profiles that were predicted from the natural pan-reactomes. The ENs were trained on the sampled panEFMs (Table S5).

This landscape also reflects a metabolite usage landscape that is based on the frequency that metabolites are used by the sampled panEFMS across random simulated environments. The metabolite usage landscape also exhibits family-specific clusters (Figure 3B) and the family-specific metabolite usage profiles that were predicted with an EN model (as explained above for the toy model and *Aeromonadaceae* pan-reactome) also behave as observed values (realizations) of the stochastic distribution covered by this evolutionary landscape (large points in Figure 3B).

The observed strong separation of prokaryotic families reflects family-specific differences in reaction content and metabolite preferences. A bias may also be expected from the way we sampled panEFMs by defining strict family-specific pan-reactome reaction pools. In reality, reaction (gene) pools are not strictly confined to a family and horizontal gene transfer between different families may alleviate the separation between family-level pan-reactomes, although we note that all reactions observed within the sequenced strains of a family were already included in the pan-reactome definition.

To get a better idea of how the reaction frequency of panEFMS sampled in random virtual environments compare to the reaction frequencies in natural pan-reactomes, we binned reactions according to their frequency in panEFMs, and plotted the distribution of these reactions in the natural pan-reactomes using violin plots (Figure S2). Although panEFMs were sampled in random virtual environments, we found that reactions with a high frequency in panEFMs are often universal among the strains of a pan-reactome (Figure S2). In contrast, there is significant variability in the frequency of reactions that are rare among panEFMs (averages usually close to 0.50, see Figure S2). Thus, our computational approach of sampling panEFMs captures at least some of the dynamics of the natural pan-genomes, that we summarized by the following scenario: (i) Reactions with high frequency in panEFMs are environment-independent (EDS = 0, Figure S3) and universally essential (Figure S2) since across many environment it is statistically unlikely for evolution to form functional reactomes without them; (ii) Reactions with intermediate frequency in panEFMs are variable in pan-reactomes and are enriched for environment-driven reactions (EDS > 0, Figure S3) since only in a fraction of the sampled environments it is statistically unlikely for evolution to form functional reactomes without them; (iii) Reactions with low frequency in panEFMs have high variability in their frequencies that are usully distributed as a U-shape (Figure S2). Their presence or absence are not captured by the panEFMs and their frequency could very well behave as the “bag of genes” models explained in the introduction (Baumdicker et al., 2012; Collins and Higgs, 2012; Haegeman and Weitz, 2012; Lobkovsky et al., 2013), although we did not explore this any further.

### Predicting metabolite-reaction associations in the pan-reactomes of 46 prokaryote families

Similar to what we observed for the toy model and for the *Aeromonadaceae* pan-reactome, most reactions have a similar predicted frequency in panEFMs across all environments (Figure S4, average Pearson r^2^ =.99; p<e-10), with some reactions exhibiting a significant environment-driven variation in frequency, quantified by the EDS and illustrated by the points that fall outside of line in Figure S4. To illustrate, we identified the two reactions that had the highest average EDS scores across all prokaryote families (Table S4): Dihydroxy hydrolase (EC 4.2.1.9) and pyruvate decarboxylase (EC 2.2.1.6). Both reactions catalyze steps in the synthesis of the three branched-chain amino acids (L-isoleucine, L-valine, and L-leucine) and are universally shared across bacterial reactomes (Amorim Franco and Blanchard, 2017). These reactions are also part of the pantothenate and coenzyme A (CoA) biosynthesis pathway, where the product of the dihydroxy hydrolase (3-Methyl1-2-oxobutanoic acid) can either be used for the synthesis of L-valine or for the synthesis of 2-Dehydropantoate, a precursor for pantothenate and subsequently CoA. Pantothenate and CoA are connected to the biosynthesis of several amino acids, which explains why reactions upstream of their synthesis would be essential or not depending on the availability of these amino-acids in the external environment.

As explained above, we used the frequency of reactions in panEFMs to train an EN model that predicts the metabolite niches of family-specific pan-reactomes from their natural reaction frequencies—these predictions are analogous to the predictions obtained from the Moran-like process that was applied to the *Aeromonadaceae* pan-reactome (Figure 2D) except that reaction frequencies are now derived from their actual distribution in the pan-reactome (Table S2) rather than from a simulated evolutionary process. In both cases the model, was trained on the frequency of panEFMs sampled across random environments. The predicted metabolic niches are summarized in Table S5. Most reactomes require inorganic ions, such as Ca, Cl, Mn, Zn, K, Mg, and Fe, and some organic molecules such as heme are also widely required (Benson and Rivera, 2013). Different pan-reactomes require specific metabolites, making them distinguishable when projected in lower-dimensional space (large points in Figure 3B). A more detailed characterization of metabolite preferences in pan-genomes can be of interest in future studies aimed at explaining the metabolic basis of genome evolution events.

### panEFMs are mechanistic predictors of patterns in pan-reactome shape, size, and distribution

To evaluate to what extent panEFMs that were sampled *in silico* from the pan-reactomes of different prokaryotic families, reflect the strains within those families, we compared multiple variables of panEFMs and pan-reactomes (Table 1). Figure 4 displays the pairwise correlations between all variables across the 46 prokaryote families, detailed in Table S6. Larger pan-reactomes contain a larger set of reactions that may be integrated into panEFMs (Pearson r = 0.94; adj. p<e-20; variable ‘pan(pEFMs)’). Notably, this variable correlates better with the size of the “shell” (Pearson r=0.85; adj. p<e-11; variable ‘shell(Reactomes)’) than with the size of the “core” (Pearson r=0.24; adj. p=0.154; variable ‘core(Reactomes)’) or the “cloud” (Pearson r=0.38; adj. p=0.016; variable ‘cloud(Reactomes)’) of the pan-reactome. We also observed a significant spread in the average size of panEFMs (Figure S5), which correlates with the average reactome size of pan-reactomes (Pearson r = 0.82; adj. p<e-10; variable ‘size(Reactomes)’) and all the other variables have similar correlations to the average panEFM size as they have with the pan-reactome size (Figure 4).

**Table 1:** Variables that were used to compare the panEFMs and pan-reactomes of 46 prokaryote families (Figure 4, Table S6).

| VARIABLE | DESCRIPTION |
| --- | --- |
| NicheBreadth | Predicted niche breadth from global environmental sequencing datasets. |
| diversity(panEFMs) | Diversity between reaction frequency of panEFMs sampled in different virtual environments (Average squared pairwise Euclidean distance) |
| fluidity(panEFMs) | Average dissimilarity between panEFMs independently of the random environments in which it was sampled |
| pan(panEFMs) | Total reactions that are included in at least one of the panEFMs sampled in different virtual environments |
| pan(Reactomes) | Number of reactions found in the pan-reactome of a prokaryote family |
| size(panEFMs) | Average size of panEFMs sampled in different virtual environments |
| size(Reactomes) | Average size of the natural reactomes from a prokaryote family |
| core(panEFMs) | Number of reactions that are present in at least 98% of the panEFMs sampled in different virtual environments |
| core(Reactomes) | Number of reactions present in at least 98% of the natural reactomes from a prokaryote family |
| shell(panEFMs) | Number of reactions that are present in 3 to 98% of all the panEFMs sampled in different virtual environments |
| shell(Reactomes) | Number of reactions that are present in 3 to 98% of the natural reactomes from a prokaryote family |
| cloud(panEFMs) | Number of reactions present in up to 3% of the panEFMs sampled in different virtual environments |
| cloud(Reactomes) | Number of reactions present in up to 3% of the natural reactomes from a prokaryote family |
| diversity(Metabs) | Diversity between metabolite usage profiles of panEFMs sampled in different virtual environments (Average squared pairwise Euclidean distance) |
| EnvDReacs | Number of reactions with an environment-driven score (EDS) EDS significantly > 0 (adj. p < 0.05 on a Z-test) |
| EnvDMetabs | Number of metabolites that are significantly associated to reactions with a non-zero EDS |

**Figure 4:**
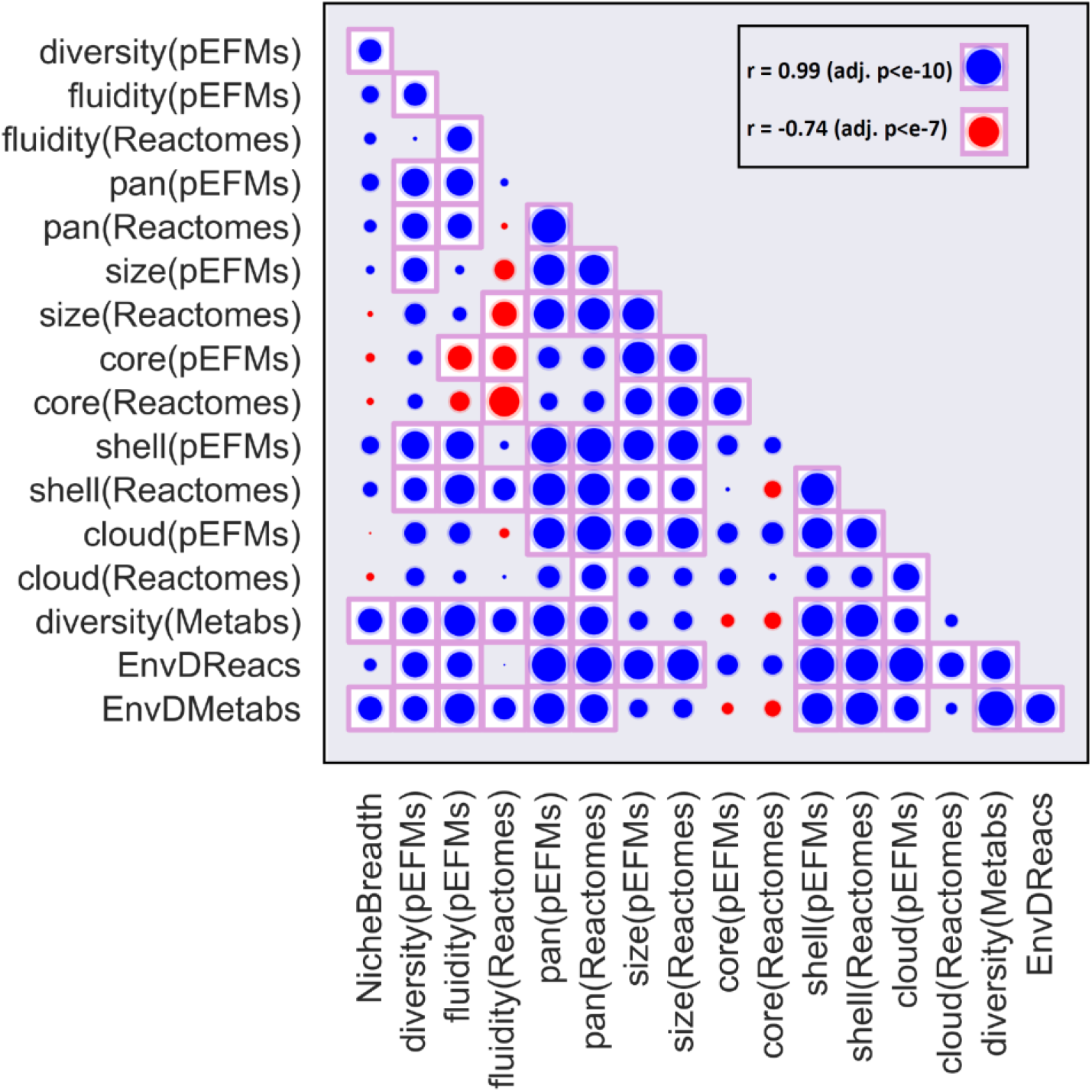
Correlation of the variables measured from panEFMs with reactomes and metagenomes across 46 prokaryotic families. A description of the variables is available in Table 1. Detailed Pearson correlation values and adjusted p-values are available in Table S6. For comparison, values of the highest (blue) and lowest (red) correlations are indicated in the top right corner.

### Correlation of panEFMs with global environmental sequencing data

Next, we evaluated whether the panEFMs and pan-reactome variables correlates with the niche breadth of prokaryote families inferred from global environmental sequencing datasets (Figure 4; Table 1; variable ‘NicheBreadth’). We inferred niche breadth scores for all families from thousands of metagenomic datasets that quantify the diversity of the environments where each family is found, and are a proxy for the breadth of their niche preferences across the planet (Von Meijenfeldt et al., manuscript in preparation, see Methods section “Niche breadth”). First, we found that the diversity between the reaction frequencies of panEFMs that were sampled from different environments positively and significantly correlated with the niche breadth of the bacterial and archaeal families (Pearson r=0.42; adj. p<e-02; variable ‘diversity(pEFMs)’). This confirmed that families whose strains occur in diverse environments tend to have more diverse environment-driven reactions than families whose strains occur in uniform environments. Notably, families with a high panEFM fluidity, that is the pan-genome analog to mutation rates (Kislyuk et al., 2011) and measures the average dissimilarity in the reaction content between random pairs of reactomes across all environments, did not have a significantly higher niche breadth (Pearson r=0.23; adj. p=0.18; variable ‘fluidity(pEFMs)’). This shows that families whose pan-reactome is capable of differentially adapting to different simulated environments may be observed in more diverse metagenomic data sets than families whose pan-reactome adapts to different environments in a similar way. The two other variables that significantly correlated with the niche breadth, namely (i) the number of metabolites that are significantly associated with environment driven reactions (reactions with and EDS significantly > 0, Pearson r = 0.46, adj. p=0.003; variable ‘EnvDMetabs’) and (ii) the diversity of metabolite usage profiles between panEFMs sampled in different environments (Pearson r=0.47; adj. p<e-03; variable ‘diversity(Metabs)’) further indicate that reactomes of families with larger niche breadths not only have more diverse environment-driven reactions, but can also use more diverse metabolite compositions when constrained to diverse environments.

### General pan-reactome patterns

By evaluating the correlation of different measures across reactomes and the collection of panEFMs sampled across 1000 virtual environments (Table 1 and Table S6) we propose a general scenario for pan-reactome evolution. For this scenario we consider that prokaryotic clades, such as the families described here, have access to a shared pool of reactions and that non-essential reactions are frequently lost. Some pan-reactomes have a large number of core reactions and also a large number of reactions that are present in all panEFMs (Pearson r=0.62; adj. p<e-04; variable ‘core(pEFMs)’). The larger the core-reactome, the less diverse are reactome pairs, measured by the fluidity (Pearson r=−0.74; adj. p<e-07; variable ‘fluidity(pEFMs)’). The shell of panEFMs and the shell of pan-reactomes are significant predictors of ecological flexibility. Large shells imply larger pan-reactomes (Pearson r=0.99; adj. p<e-46), more fluidity (Pearson r=0.63; adj. p<e-04), and more environment-driven reactions (Pearson r=0.94; adj. p<e-21) and metabolites (Pearson r=0.77; adj. p<e-08). These scenarios are non-trivial properties of the pan-reactome composition, mechanistically identifiable in the collection of panEFMs and suggest that the shape and size of the pangenomes reflect evolution in different environments.

## Discussion

Diverse evolutionary groups have evolved specific sets of metabolic reactions to obtain their required energy and biomass. On average, more than 50% of the genes in microbial genomes code for metabolic functions (Griesemer et al., 2018) and metabolic genes are often found to be horizontally transferred (Goyal, 2018). The different sets of metabolic reactions used in different contexts by microbes reflect patterns and mechanisms of their genome evolution. For this reason, reactomes have increasingly been used as model systems for evolutionary genomics (Aguilar-Rodríguez and Wagner, 2018; Barve et al., 2012; Pál et al., 2006; Pang and Lercher, 2019; Szappanos et al., 2016; Wagner, 2009). Here we modelled the evolution of reaction frequency distribution in pan-reactomes to understand the forces and mechanisms driving genome evolution. We used pan-reactomes as proxies for pan-genomes because they represent complex genotype-to-phenotype maps that allow us to directly explore the effects of differential gene composition, and study the composition of genomes in the context of complex cellular phenotypes.

We developed a mechanistic evolutionary model to expose the forces that drive reaction frequency distribution. In our framework, pan-reactomes share a pool of reactions and individual reactome lineages undergo a process of extensive gene loss. We modeled this natural process of gene loss (Bolotin and Hershberg, 2016; Puigbò et al., 2014; Snel et al., 2002; Wolf and Koonin, 2013), by allowing reactomes to lose all of their non-essential reactions in independent iterations across different simulated environments. This process provided an empirical distribution of the functional reactomes that can evolve from a given reaction pool. This distribution allowed us to disentangle the environment-driven (nutrition) and environment-independent (nature) reactions, build a model that can predict metabolic niches from reaction frequencies, and compare reactome patterns between prokaryote families.

Previous studies have also used stochastic reductive evolution in reactomes to assess alternative scenarios of reactome diversity (Barve et al., 2012; Szappanos et al., 2016; Wagner, 2009). Most of these studies were applied to the pan-reactome of the *Escherichia coli* clade (Pál et al., 2006; Szappanos et al., 2016; Yizhak et al., 2011) or were applied to understand patterns that emerge from the universal set of reactions (Barve et al., 2012; Wagner, 2009), i.e. all metabolic reactions that have been identified in prokaryotes. Here we applied stochastic reductive evolution to understand differences within and across pan-reactomes of different prokaryotic families, providing a unique systematic overview of their pan-reactome dynamics. This approach allowed us to expose the patterns in pan-reactome size, shape, and diversity that are functions of its composition. It also allowed us to predict family-specific metabolic niches that await experimental testing.

We identified patterns at two levels. At the lower level of individual reactions, we mechanistically predicted the essentiality of a reaction based on the capacity of the pan-reactome to generate functional alternatives across environments. This allowed us to identify how many and which reactions become essential in new environments. At the higher level of pan-reactomes, our framework revealed non-trivial features. While all 46 pan-reactomes were subjected to the same process of reaction loss and were under similar functional constraints, features such as size, shape, and diversity were significantly different between families. We found that these features depend closely on the composition of the pan-reactomes and are reflected in the *in silico* generated collection of panEFMs.

The composition of the pan-reactome determines patterns observed in specific organisms. With our framework, we mechanistically identified these patterns from functional reactomes. For example, some pan-reactomes form functional reactomes with a large number of core reactions. These reactomes are very similar to each other and use a small set of metabolites (Figure 4, Table S6). Other pan-reactomes form functional reactomes with a small set of core reactions and a large set of reactions of intermediate frequency (shell reactions). These require more metabolites and exhibit significantly different reaction frequencies when their pan-reactomes are challenged with different environments (Figure 4, Table S6). All these properties result from the different ways that reaction sets can assemble to form functional reactomes.

Reaction frequency highly depends on the global reactome functionality since the patterns that we observed in panEFMs were identified without determining specific evolutionary goals or additive adaptive values for specific reactions. This constraint of functionality shapes the sample space of possible reactomes and constraints its evolutionary potential. In a similar trend, we expect that the combinatorial functionality of genes within a gene pool is an important driver of the pan-genome composition. In other words, the question of how often a gene is found in the genomes of a prokaryote group is to some extent addressed by how often the gene is expected in functional gene sets and to some extent on the composition of its external environment. We used these connections to predict the metabolic niches of natural evolving pan-genomes and identify the forces that shape these pan-genomes as important functional units of prokaryote evolution.

## Methods

### Reactomes

Bacterial and archaeal strains (n = 4885) from 46 taxonomic families were selected from the PATRIC database (Wattam et al., 2017) (Table S2). We chose to use families that had genomes sequences of over 24 different species and selected one strain of each species based on the maximum completeness and minimum contamination values of their genome sequences as reported in the PATRIC metadata.

We reconstructed genome-scale metabolic models (GSMMs) for each strain using the model SEED pipeline (Henry et al., 2010) implemented in PATRIC with the Mackinac python package (Mundy et al., 2017). Each model contains a list of reactions that are predicted to be coded by the prokaryote genomes; these reactions are referred to as the reactome of the strain. In addition to the genome encoded reactions, each model has a biomass reaction consisting of the relative proportions of biomass components, such as amino acids, nucleotides, proteins, fats, co-factors, and sugars, that the reactome should be able to synthesize in a growth environment. Additionally, some reactions that were not annotated in the genomes were added to assure that the reactomes were capable of producing biomass in complete media, an approach referred to as “gap-filling” (Pan and Reed, 2018).

The functionality of GSMMs was assessed by flux balance analysis (FBA) (Feist and Palsson, 2010), optimized for biomass production. We used the flux yields on the biomass reaction as an indication of growth.

### Pan-reactomes

Pan-reactomes were generated by merging the reactomes of all the strains from a given prokaryote family. Each reaction was added once. Additionally, we added exchange reactions for the compounds that have transporters in any of the reactomes (Table S2), resulting in pan-reactomes with the same group of 292 exchange reactions.

### Environment ball

We generated vectors of random uniform relative concentrations for the shared list of external compounds added as exchange reactions, excluding water and oxygen (n=290). For obtaining relative uniform concentrations, we sampled a Dirichlet distribution with dimensions equal to the number of compounds (290) with uniform parameters). Samples from this distribution add to one and there is an equal probability of observing any relative concentration of any of the compounds. The resulting relative concentrations were adjusted to a constant uptake rate of water in mmol.gDW^−1^.h^−1^. Oxygen was added as a binary factor, with environments being either aerobic (containing an unconstrained amount of oxygen) or anaerobic (with zero oxygen), selected with a probability of 0.5. We generated 1000 random samples of the environment ball and used the resulting concentrations as the growth environments (Table S2).

### Toy model

The toy model was generated with the reactions in Table S1. Functionality was directly assessed by evaluating if the biomass components could be synthesized without accumulating by-products. Since stoichiometries were all equal to one (Table S1, Figure 1), the external environment was defined by the presence or absence of metabolites.

#### Moran Process

We evolved populations of reactomes derived from the toy and the *Aeromonadaceae* pan-reactomes with a Moran-like process (Moran, 1958). For this, we started with random functional reactomes (n=1000) and simulated a two-step process. In the first step, a random reactome was chosen and a reaction was either deleted or inserted. If the change resulted in a functional reactome, the process continued, otherwise, the previous reactome was restored. In the second step, two reactomes were chosen and one of them was replaced by a copy of the other, simulating a birth-death process with a constant population size. After many iterations (n=10^6^), the different types of reactomes that persisted were selected as the evolved reactome types.

#### Sampling the environment-specific collection pan-reactome elementary flux mode (panEFMs)

To generate random samples from the environment-specific collection of panEFMs, we first constrained the pan-reactome to a given environment (Table S2) and only proceeded if the flux on the biomass reaction was greater than zero with five significant digits. We then randomly removed reactions from the pan-reactomes and evaluated if the resulting network exhibits flux in the biomass reaction that is greater than a cutoff of 1% of the flux observed in the pan-reactome. If the biomass flux is below the cutoff, the reaction was restored to the network, otherwise, the next reaction was removed until all reactions were assessed. At the end of one iteration, the reactions that remained in the network constitute a random sample of a panEFM. Next, we randomized the order of reaction removal and repeated the process. Each randomization of the reaction order finds a random panEFMs. We sampled 1000 panEFMs for each of the 1000 metabolite concentrations in the environment ball.

#### Environment-driven scores (EDS)

Reaction frequency in the environment-specific collections of panEFM was represented by a matrix containing the different environments of the environment ball as rows and reactions found in a pan-reactome as columns. Similarly, environment-specific metabolic niches were summarized in a matrix with a similar structure but containing the environment ball metabolites as columns. The averages of the columns of these matrices are, respectively, the expected values of reaction frequencies and the expected values of metabolite usage across environments. Residuals were obtained by taking the difference between these expected values and the values in each row (each environment). The environment-driven scores for reactions and metabolites were defined as the standard deviation of these residuals, divided by their maximum value.

#### Obtaining metabolite usage profiles

The metabolic niches of the panEFMs of bacterial and archaeal families were obtained by enumerating which of the possible external compounds were imported into the metabolic network when optimizing for biomass production. For each panEFM we first obtained the environment-specific FBA solution. We then assessed the fluxes in the exchange reactions for this solution. Negative fluxes correspond to the metabolites that are effectively required to produce biomass. One metabolic niche corresponds to the set of metabolites whose transporters exhibited a negative flux in the FBA solution of a panEFM. Metabolic niches were summarized by the metabolite frequencies obtained from the 1000 random samples of panEFMs that were obtained for each environment. We thus obtained a metabolic niche for each random environment by enumerating how often each metabolite was used after sampling panEFMs from pan-reactomes.

#### Reaction frequencies

We restricted our analysis to reactions that had gene evidence (not gap-filled) and that could be active in a model. To define if a reaction could be active, we used flux variability analysis and excluded reactions that exhibited a flux variability of zero. We refer to the “natural reaction frequencies” as the frequencies that were observed from the reaction composition of reactomes reconstructed from the genomes of a prokaryote family, while “frequencies in panEFMs” refer to the frequency that reactions were found in the random samples of panEFMs.

#### Elastic net

An elastic net model was trained to predict metabolite usage from reaction frequencies (natural or resulting from simulations in the Moran process). We used the matrices described above as training sets with five-fold cross-validation. The natural reaction frequencies were used to predict evolutionary environments. To train and fit the model we used the Python 3.7 package scikit-learn version 0.22.2.

#### Niche breadth

For each family we calculated its niche breadth on the scale from specialist to generalist based on its presence in a large number of publicly available environmental sequencing datasets. This measure will be presented elsewhere in detail (Von Meijenfeldt et al., manuscript in preparation) but we will discuss the reasoning here.

In short, we selected taxonomically annotated environmental sequencing projects from the MGnify dataset(Mitchell et al., 2020). We selected analyses that were annotated with the 4.1 pipeline to ensure that the taxonomic profiles were comparable, removed analyses with less than 50,000 taxonomically annotated reads or >= 10% eukaryotic reads, chose a maximum of 1,000 samples per biome, and selected 1 analysis per sample. The 22,518 selected analyses spanned 140 different biomes across a wide geographical range, containing both metagenomic, transcriptomic, and amplicon datasets.

A family was considered present in a sample if its relative abundance was >> 1/10,000. Niche breadth was defined as the mean pairwise distance between all the samples in which a family is found, where the mean pairwise distance is defined as 1/2 - (Spearman’s rank correlation on family level / 2). Since this measure is solely based on the taxonomic content of a sample, it is independent of manually added metadata such as the biome from which it originates. A family with a low score is primarily found in samples with similar taxonomic profiles and we thus consider a specialist, and a family with a high score is found in more dissimilar samples and is thus a generalist.

## Acknowledgments

DRG was supported by CNPq Science Without Borders. FABvM and BED were supported by the Netherlands Organization for Scientific Research (NWO) Vidi grant 864.14.004 and European Research Council (ERC) Consolidator grant 865694: DiversiPHI.

## Supplementary Figures

**Figure S1:**
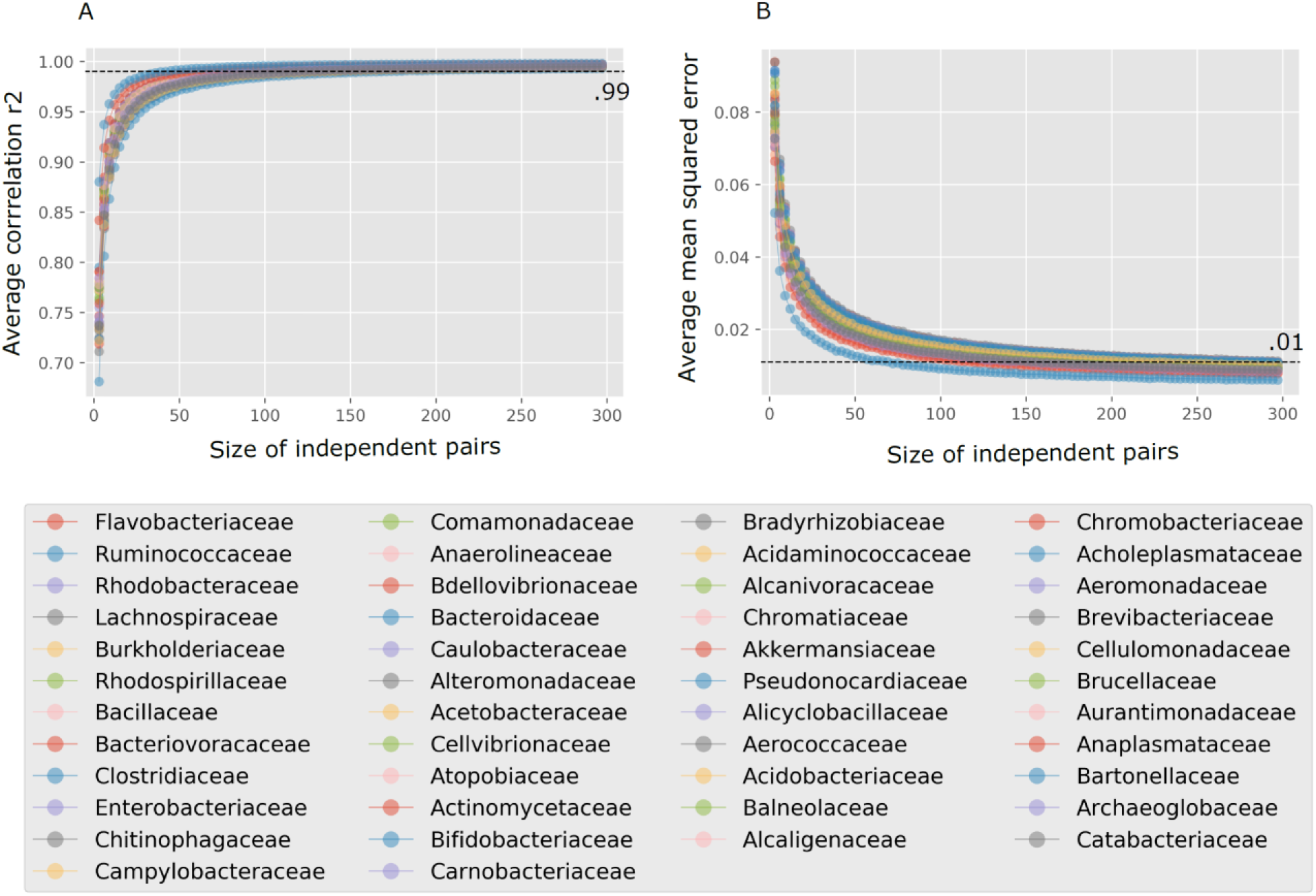
Convergence of the reaction frequencies of panEFMs sampled across 1000 virtual environments (Table S2) to an average. The frequency of reactions was obtained from two random-independent (non-overlapping) groups of panEFMs defined across the environment ball. The group sizes varied according to the x-axis. As a measure of convergence, we used the correlation (A) and the mean-squared error (B) between the frequencies of two independent groups. Within a prokaryote family, these values were averaged for the 1000 different environments. Dashed lines show the .99 and .01 correlation and mean-squared error values, respectively.

**Figure S2:** Distribution of pan-reactome reaction frequencies and panEFMs reaction frequencies. The frequency in panEFMS are binned in a frequency gradient and the violin plot show the frequency distribution of the reactions within a bin in the natural pan-reactomes.

Accessible at: https://github.com/danielriosgarza/NutritionOrNature/blob/main/FigureS2.pdf

**Figure S3:** Distribution of the environment-driven reaction score (EDS_ and panEFMs reaction frequencies. The frequency in panEFMS are binned in a frequency gradient and the violin plot show the distribution of the EDS computed for the reactions within a bin.

Accessible at: https://github.com/danielriosgarza/NutritionOrNature/blob/main/FigureS3.pdf

**Figure S4:**
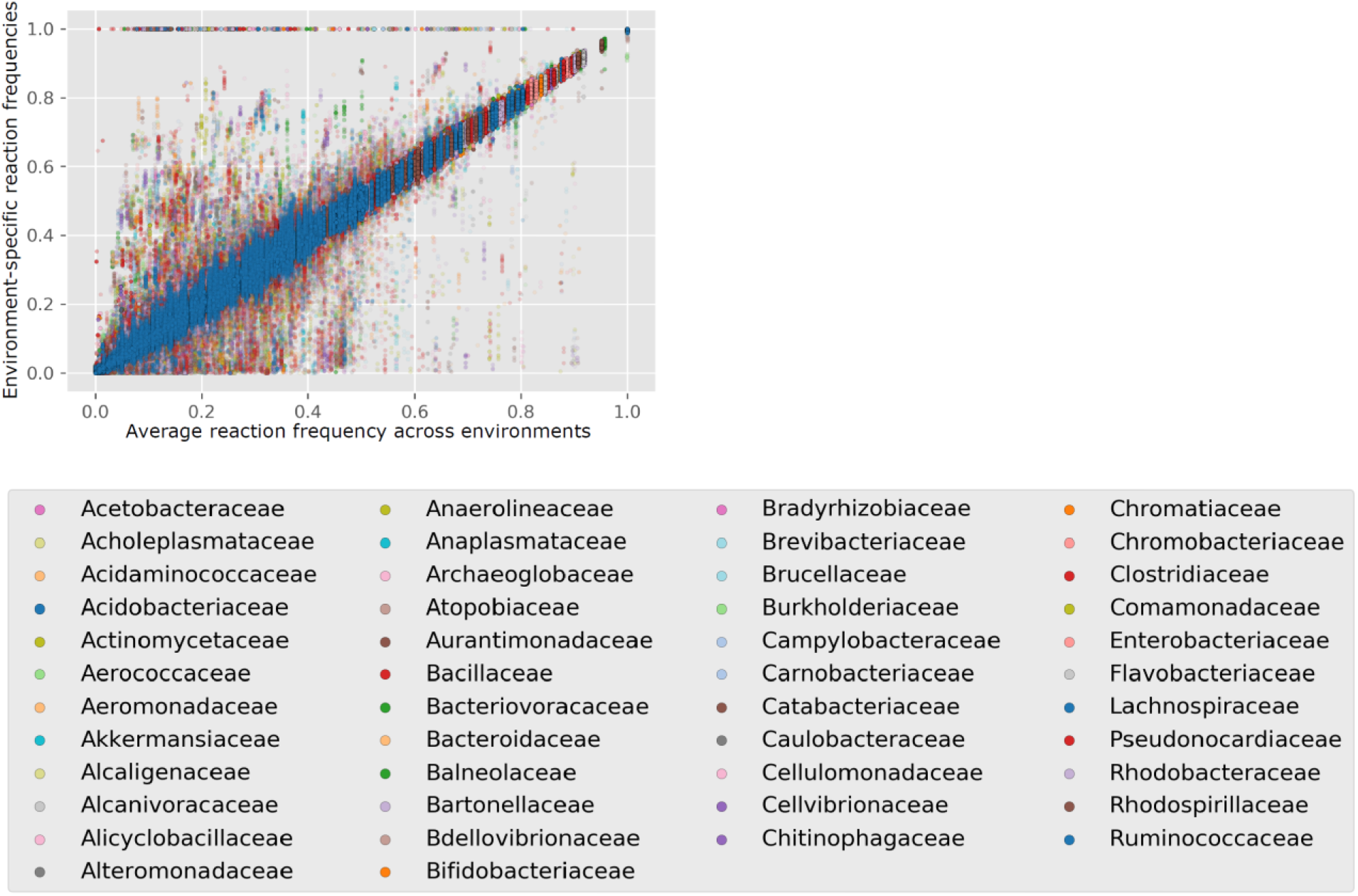
Comparison of panEFMs across environments. Scatter plot of the average reaction frequency of panEFMs defined across random virtual environments and within each environment.

**Figure S5.**
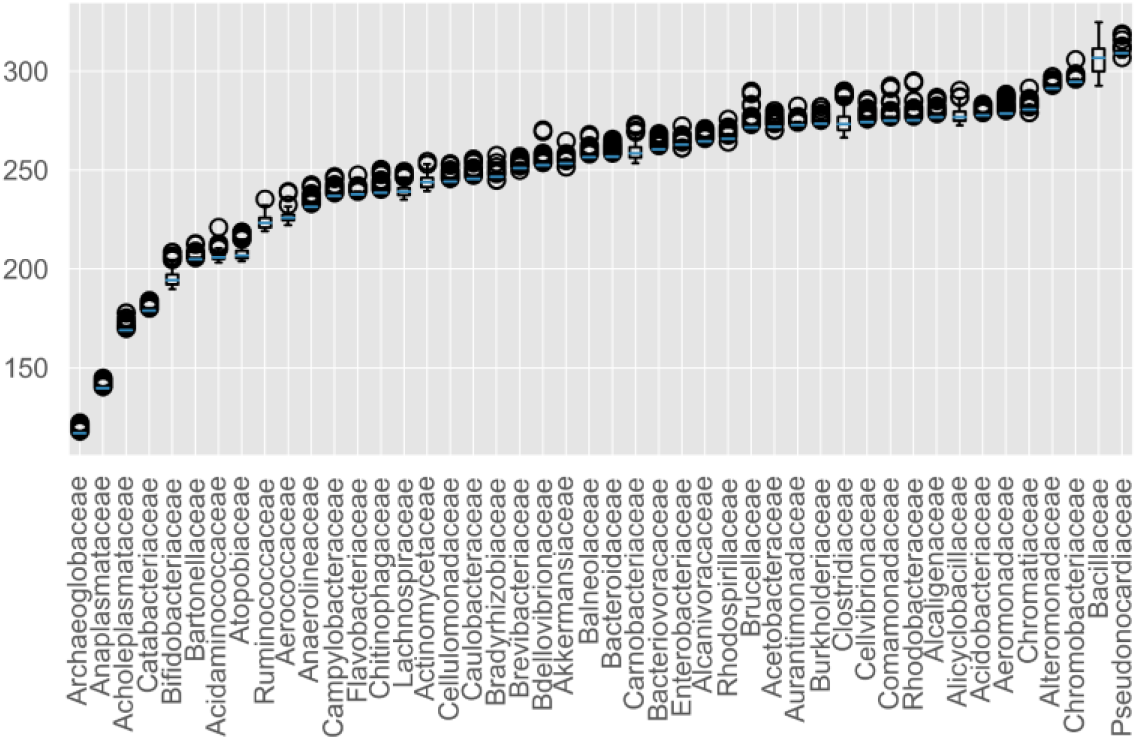
Size distribution of panEFMs sampled across random virtual environments.

## Supplementary Tables

**Table S1:** Reactions in the toy model. List of toy reactions used in the *in silico* simulations of the toy model displayed in Figure 1.

| Reaction | Equation |
| --- | --- |
| R1 | $M1_e \rightarrow M1_i + M2_i$ |
| R2 | $M2_e \rightarrow M1_i + M3_i$ |
| R3 | $M3_e \rightarrow M4_i$ |
| R4 | $M4_e \rightarrow M5_i$ |
| R5 | $M5_e \rightarrow M6_i$ |
| R6 | $M6_e + M7_e \rightarrow M7_i$ |
| R7 | $M8_e \rightarrow M8_i$ |
| R8 | $M9_e \rightarrow M8_i$ |
| R9 | $M1_i \rightarrow M9_1$ |
| R10 | $M2_i \rightarrow M10_e$ |
| R11 | $M7_i \rightarrow M8_i$ |
| R12 | $M8_i + M6_i \rightarrow M10_i$ |
| R13 | $M5_i \rightarrow M11_i$ |
| R14 | $M3_i + M9_i \rightarrow M12_i$ |
| R15 | $M4_i \rightarrow M5_i + M3_i$ |
| Biomass | $M10_i + M11_i + M12_{12}$ |

**Table S2:** Bacterial and archaeal strains used in this study. The Aeromonadaceae strains are highlighted.

Accessible at: - https://github.com/danielriosgarza/NutritionOrNature/blob/main/Table_S2.xlsx

**Table S3:** Environment compounds used in this study. Concentrations of 1000 random environments that constitute the environment ball (see Methods). These concentrations are set as upper bounds to the metabolic models of the pan-reactomes.

Accessible at: https://github.com/danielriosgarza/NutritionOrNature/blob/main/Table_S3.xlsx

**Table S4:** Environment-driven reaction scores (EDS) for all 46 prokaryote families.

Accessible at: https://github.com/danielriosgarza/NutritionOrNature/blob/main/Table_S4.xlsx

**Table S5:** Elastic net predictions of the metabolite usage by the pan-reactomes of 46 prokaryote families

Accessible at: https://github.com/danielriosgarza/NutritionOrNature/blob/main/Table_S5.xlsx

**Table S6:** Correlation of variables related to panEFM, pan-reactomes, and metagenomes

Accessible at: https://github.com/danielriosgarza/NutritionOrNature/blob/main/Table_S5.xlsx

